# Environmental Oxygen Regulates Astrocyte Proliferation to Guide Angiogenesis during Retinal Development

**DOI:** 10.1101/861948

**Authors:** Robin M Perelli, Matthew L O’Sullivan, Samantha Zarnick, Jeremy N Kay

## Abstract

Angiogenesis in the developing mammalian retina requires patterning cues from astrocytes. Developmental disorders of retinal vasculature, such as retinopathy of prematurity (ROP), involve arrest or mispatterning of angiogenesis. Whether these vascular pathologies involve astrocyte dysfunction remains untested. Here we demonstrate that the major risk factor for ROP – transient neonatal exposure to hyperoxia – disrupts formation of the angiogenic astrocyte template. Exposing mice to hyperoxia (75% O_2_) from postnatal day (P)0-4 suppressed astrocyte proliferation, while return to room air (21% O_2_) at P4 triggered extensive proliferation, massively increasing astrocyte numbers and disturbing their spatial patterning prior to arrival of developing vasculature. Proliferation required astrocytic HIF2α and was also stimulated by direct hypoxia (10% O_2_), suggesting that astrocyte oxygen sensing regulates the number of astrocytes produced during development. Along with astrocyte defects, return to room air also caused vascular defects reminiscent of ROP. Strikingly, these vascular phenotypes were more severe in animals that had larger numbers of excess astrocytes. Together, our findings suggest that fluctuations in environmental oxygen dysregulate molecular pathways controlling astrocyte proliferation, thereby generating excess astrocytes that interfere with retinal angiogenesis.

## Introduction

Precise coordination between growing neurons, glia, and blood vessels is a central feature of neural development and is essential for building a functional nervous system. A striking example of coordinated neuro-glial-vascular growth occurs at the vitreal surface of the mammalian retina within the retinal nerve fiber layer (RNFL). This layer contains several different structures that are essential for visual function, including the intrinsic retinal vasculature; the retinal ganglion cell (RGC) axons, which course through the RNFL towards the optic nerve head; and a population of glia known as RNFL astrocytes. During development, astrocytes and blood vessels enter the retina at the optic nerve head and spread centrifugally through the RNFL to colonize the entire retinal surface (Selvam et al., 2018). These migratory events are coordinated through a sequential series of cell-cell interactions: RGC axons guide astrocyte migration, and astrocytes in turn guide endothelial cell growth during angiogenesis (Dorrell et al., 2002; Fruttiger et al., 1996; Gerhardt et al., 2003; O’Sullivan et al., 2017). When this sequence of events is disrupted there are major deleterious consequences for development of vasculature, including severe disruption or even arrest of angiogenesis (Duan and Fong, 2019; Fruttiger et al., 1996; O’Sullivan et al., 2017; Tao and Zhang, 2016). Furthermore, the delay or arrest of RNFL angiogenesis is a key hallmark of human retinal developmental disorders such as retinopathy of prematurity (ROP) (Foos, 1987; Hellström et al., 2013). It is therefore of great interest to understand the developmental mechanisms that enable the orderly progression of retinal angiogenesis and how these mechanisms might become perturbed in the context of ROP pathology.

The principal risk factors for ROP are prematurity, low birth weight, and supplemental oxygen. In infants affected by ROP, halted extension of nascent vessels leaves the peripheral retina avascular (Foos, 1987). This is commonly termed Phase I of ROP. Subsequently, in Phase II, hypoxia in the ischemic peripheral retina stimulates uncontrolled neovascular growth, which can lead to retinal detachment and ultimately vision loss (Hellström et al., 2013). Mechanistically, exposure to oxygen is thought to provoke ROP because hyperoxia diminishes the molecular drive for angiogenesis. As such, hyperoxia could explain the wavefront arrest. However, it remains unclear why vessels do not resume their orderly progression through the RNFL once normoxic conditions are restored.

One possible explanation for this failure of wavefront progression is that the astrocytic template for angiogenesis becomes disturbed. In normal development, astrocytes arrange their somata and arbors into a pre-pattern resembling a capillary bed. This astrocyte template exerts powerful effects on the ultimate pattern of the vascular network, and is required for normal vascular morphogenesis (Fruttiger et al., 1996; Gerhardt et al., 2003; O’Sullivan et al., 2017; Tao and Zhang, 2016). Despite this well-established role of astrocytes in RNFL vascular development, the role astrocytes play in ROP remains obscure. As in mice, human astrocytes also migrate ahead of the vasculature where they are positioned to influence angiogenesis (Chan-Ling et al., 2004). Furthermore, ROP histopathological studies have noted that the fibrovascular ridge – a pathological structure that forms at the junction between vascular central and avascular peripheral retina – is composed predominately of astrocytes, whereas few astrocytes are detected peripheral to this ridge (Sun et al., 2010). Thus, there is reason to think that astrocyte patterning might be disturbed in ROP. However, despite some hints from animal studies (Duan et al., 2017; Morita et al., 2016; Zhang et al., 1999), it is presently unknown whether astrocyte development is impacted by ROP risk factors such as hyperoxia. Resolving this issue has the potential to reveal new developmental mechanisms regulating normal astrocyte development, and would clarify whether altered astrocyte development is a plausible candidate to underlie ROP vascular pathology.

To investigate these questions, we sought to understand astrocyte development in a mouse model of oxygen-induced retinopathy (OIR). Since the identification of oxygen as the major modifiable risk factor for ROP, several animal OIR models have been developed. In the most common model, mouse pups are exposed to 75% O_2_ from postnatal day 7 (P7) to P12 (Smith et al., 1994). This manipulation causes central capillary vaso-obliteration during the period of hyperoxia followed by neovascularization upon return to room air. This paradigm has proven to be a powerful tool for exploring molecular mechanisms of neovascularization and oxygen toxicity. However, it has some limitations, particularly as a model of Phase I ROP. First, the phenotype of mouse OIR – central vaso-obliteration and neovascularization – does not model the peripheral avascularity seen in ROP, nor the hyperplasia at the vascular-avascular junction (Foos, 1987). Second, the P7 onset of oxygen manipulation is after the majority of astrocyte development is complete and primary vascular plexus has been established, which places the physiologic insult at a later ontogenetic stage than ROP. For these reasons, we explored the possibility of starting hyperoxia exposure at earlier stages. Unlike in standard OIR, a period of hyperoxia at P0 can cause long-term retinal vascular pathology with features reminiscent of advanced ROP (Lajko et al., 2016; McMenamin et al., 2016). Thus, mouse models featuring early hyperoxia may have utility in understanding Phase I of ROP and the involvement of astrocytes. However, other studies have reported minimal long-lasting vascular effects following P0 hyperoxia (Morita et al., 2016), so the extent to which this paradigm serves as a useful disease model remains uncertain.

Here we show that neonatal exposure to hyperoxia has long-lasting consequences for development of RNFL astrocytes and vasculature. In our neonatal oxygen-induced retinopathy (NOIR) paradigm, mice were raised in hyperoxic conditions starting at P0 and returned to room air at P4. Retinal defects including vitreous hemorrhage and retinal degenerative changes were found to persist for weeks after return to normoxia. To understand the origin of these defects we examined development of vasculature and astrocytes in this model. Consistent with previous reports (Morita et al., 2016; West et al., 2005), we found that the hyperoxic phase of the protocol prevents retinal angiogenesis while permitting astrocyte migration. Return to normoxia triggered a surge in astrocyte mitotic activity producing a vast excess of astrocytes at an age when the astrocyte population normally dwindles (Puñal et al., 2019). Environmental hypoxia similarly stimulates astrocyte proliferation, suggesting that a relative decrease in oxygen levels is the mitogenic stimulus. Astrocyte proliferation following hyperoxia was blunted by cell-type specific knockout of HIF2α, indicating that oxygen-sensitive transcriptional programs intrinsic to astrocytes regulate their mitotic activity. Following the initial proliferative response, the severity of persistent astrocyte defects was variable between animals. Strikingly, the severity of vascular defects was strongly correlated with the severity of the excess astrocyte phenotype. We propose a model in which the number of retinal astrocytes is modulated by oxygen through HIF2α, and that dysregulation of this pathway perturbs formation of the angiogenic astrocyte template leading to defective angiogenesis.

## Methods

### Mice

All experiments were conducted with the approval and under the supervision of the Duke University IACUC. For experiments with wild type mice, timed pregnant CD-1 mice were purchased from Charles River (Wilmington, MA). *GFAP-Cre* mice, with the human GFAP promoter driving expression of Cre recombinase (Zhuo et al., 2001), were acquired from Jackson Laboratory (Jax stock 004600). HIF2α-flox (*Epas1*^*tm1Mcs*^) mice (Gruber et al., 2007) were also acquired from Jackson Laboratory (Jax stock 008407). For experiments with CD-1 mice, an experimental cohort of 2-4 timed-pregnant animals were monitored at ∼8 hour intervals to identify the time of birth. Mothers that did not deliver within 8 h of the rest of the cohort were removed from the experiment. For the remaining litters, pups were randomly assorted and cross-fostered amongst the dams in the experimental cohort to control for effects of litter and minor variation in birth timing. For each experiment, the number of pups per cage was matched between cages assigned to normoxic and experimental groups. Across all experiments the number of pups per cage was between 8 and 12. Animals of both sexes were used for all experiments.

### Environmental oxygen manipulation

Cages with litters of mice and their mothers were placed inside an environmental chamber (A15274P, BioSpherix) to regulate oxygen concentrations. Medical O_2_ or N_2_ was mixed with room air by a regulator and O_2_ concentration calibrated and monitored with an O_2_ sensor (ProOx Model 360, BioSpherix). 75% O_2_ was used for hyperoxia and 10% O_2_ for hypoxia.

### Immunohistochemistry

Mice were anesthetized with ice or isoflurane, rapidly decapitated, eyes removed and immersion fixed in 4% paraformaldehyde for 1.5 hours at 4°C before being stored in phosphate buffered saline (PBS) at 4°C. For flat-mounts, retinas were dissected free from the fixed eyes and blocked at room temperature for 1 hour in PBS with 0.03% Triton X-100 (Sigma-Aldrich) and 3% Normal Donkey Serum (Jackson Immunoresearch). Retinas were then stained with primary antibody for 5-7 days at 4°C, washed 3 times with PBS, and then stained with donkey secondary antibodies (Jackson Immunoresearch) at a standard dilution of 1:1,000. Following immunostaining, 4 relieving cuts were made in the retinas and they were flat-mounted on cellulose membranes (Millipore HABG01300) on glass slides and coverslipped with Fluoromount-G (Southern Biotech).

For cryosections, fixed whole eyes were sunk in 30% sucrose in PBS overnight. 20 μm sections were then cut using a cryostat. Sections were hydrated for 10 minutes with PBS and blocked for 30 minutes in PBS with 0.03% Triton X-100 (Sigma-Aldrich) and 3% Normal Donkey Serum (Jackson Immunoresearch). Sections were incubated in primary antibodies overnight, washed three times with PBS, stained with secondary antibodies for two hours, and washed twice with PBS before mounting with Fluoromount-G.

Primary antibodies used were as follows: goat anti-GFAP (1:1,000, Abcam ab53554); rat anti-Ki67 (1:3,000, Ebioscience 14-5698-80); mouse anti-neurofilament (1:1,000, EMD Millipore MAB1621); rabbit anti-Pax2 (1:200, Covance PRB-276P); rat anti-PDGFRα (1:1000, BD Biosciences 558774); rabbit anti-Sox9 (1:4,000, Millipore AB5535); goat anti-VEGF-A (1:500, R&D Systems, AF-493-SP); Griffonia simplicifolia Isolectin B4 (IB4; 1:100, Life Technologies; Alexa 488-conjugated I21411, or biotinylated I21414) was included with primary antibodies to stain blood vessels.

### Microscopy and image analysis

Retinas were imaged on a Nikon A1R confocal laser scanning microscope with 4x air, 20x air or 60x oil immersion objective lenses. A resonant scanner and motorized stage were used to acquire whole-retina tile scan images. Images were then processed and analyzed in FIJI/ImageJ (Schindelin et al., 2012). En-face images of vasculature depict Z-projections of confocal stacks. The projected slices were selected to encompass the innermost vascular plexus to the extent possible, although in some images some deeper vasculature is also visible. Sox9 and Sox9/Ki67 double positive astrocytes were counted manually in 60x images. Total astrocyte numbers were estimated by multiplying retina area by average weighted astrocyte density; 3-4 fields of view from central, mid-peripheral, and peripheral eccentricities were analyzed, mean density for each eccentricity in each retina calculated, and overall average density calculated by weighting central, middle, and peripheral eccentricities with factors of 0.11, 0.33, and 0.56, respectively, based on a circular approximation of the retina divided into 3 zones by concentric rings with radii of 1x, 2x, and 3x (O’Sullivan et al., 2017). For hypoxia experiments, we found that rearing mice in 10% O2 inhibited migration of astrocytes into peripheral retina. Therefore, to avoid confounding our proliferation analysis, we limited our quantification of Ki67^+^ astrocytes to the central retinal region that astrocytes were able to colonize.

### Astrocyte and blood vessel coverage

To assess retinal coverage by astrocytes and blood vessels, low-magnification tile-scan images of whole retinas were used. The retinal perimeter was manually traced in ImageJ to measure the area of each retina, and within that perimeter a second curve encompassing furthest peripheral Sox9 or IB4 stained area was drawn and its area measured to yield astrocyte and blood vessel area, respectively.

### Analysis of HIF2α mutant astrocytes: VEGF-A expression and proliferation

To assess HIF pathway activation on a cell-by-cell basis, retinal whole-mounts were stained with VEGF-A, Sox9, and Ki67 antibodies as described above. Confocal image stacks were acquired at 60x. Sox9^+^ astrocytes were classified as VEGF-A^+^ or VEGF-A^−^ based on these images. For proliferation analysis at P2, the fraction of Ki67^+^ astrocytes in each category was quantified from 3 *GFAP-Cre;* Hif2α-flox mutants and 3 wild-type littermates. Data were plotted as the 95% confidence interval of the sample proportion (n = 2151 wild-type astrocytes, n = 1356 mutant astrocytes were analyzed from the 3 mice of each genotype). To assign P10 *GFAP-Cre;* Hif2α-flox mutants as either “VEGF-high” or “VEGF-low,” we assessed 20x and 60x image stacks and examined tissue by eye. With only one exception, all mutants examined could be easily assigned to one of these categories because the vast majority of the astrocyte population was either VEGF-A^+^ or VEGF-A^−^ (n = 17 mutants from 5 separate litters). In the one exception we observed a mixture of VEGF-A^+^ and VEGF-A^−^ astrocytes; this animal was excluded from the analysis.

### Data analysis

Statistical analyses were performed in GraphPad Prism 8 (GraphPad Software) or JMP Pro 13 (SAS Institute). Tests for statistical significance included two-tailed unpaired T-tests; 1-way ANOVA with post-hoc Tukey’s test; and 2-way ANOVA with post-hoc Holm-Sidak test. Post-hoc test P-values were corrected for multiple comparisons. Summary statistics reported as mean ± s.d., and error bars display s.d.

## Results

### Neonatal hyperoxia disrupts retinal vascular development

To investigate how early disruptions in oxygenation affect retinal vascular development, we raised newborn mice in 75% O_2_ or room air (21% O_2_) from P0 to P4 (Fig. 1A). Vessel development was assessed during and after the hyperoxia period (P4-P21) by staining flat-mounted retinas with the endothelial cell marker isolectin B4 (IB4), and quantifying the retinal area covered by vessels. In control mice, vessels progressed radially from central to peripheral retina, completely filling the RNFL between P8-P12 (Fig. 1B,C). By contrast, in mice exposed to P0-P4 hyperoxia, angiogenesis was significantly delayed (Fig. 1B,C). Consistent with prior reports (Morita et al., 2016; West et al., 2005), hyperoxia completely prevented blood vessel sprouting from the optic nerve head (ONH) with the rare exception of a few stray vessels escaping into the proximal portions of the retina. Only after the mice were returned to normoxic conditions did the primary vascular plexus begin to grow (Fig. 1B,C). During this phase of development, centrifugal extension of the vascular wavefront was more variable in hyperoxia-exposed mice compared to controls. Eventually, however, the vasculature was able to reach the retinal margin in all treated mice, such that the entire nerve fiber layer was vascularized by P21 (Fig 1C).

**Figure 1.**
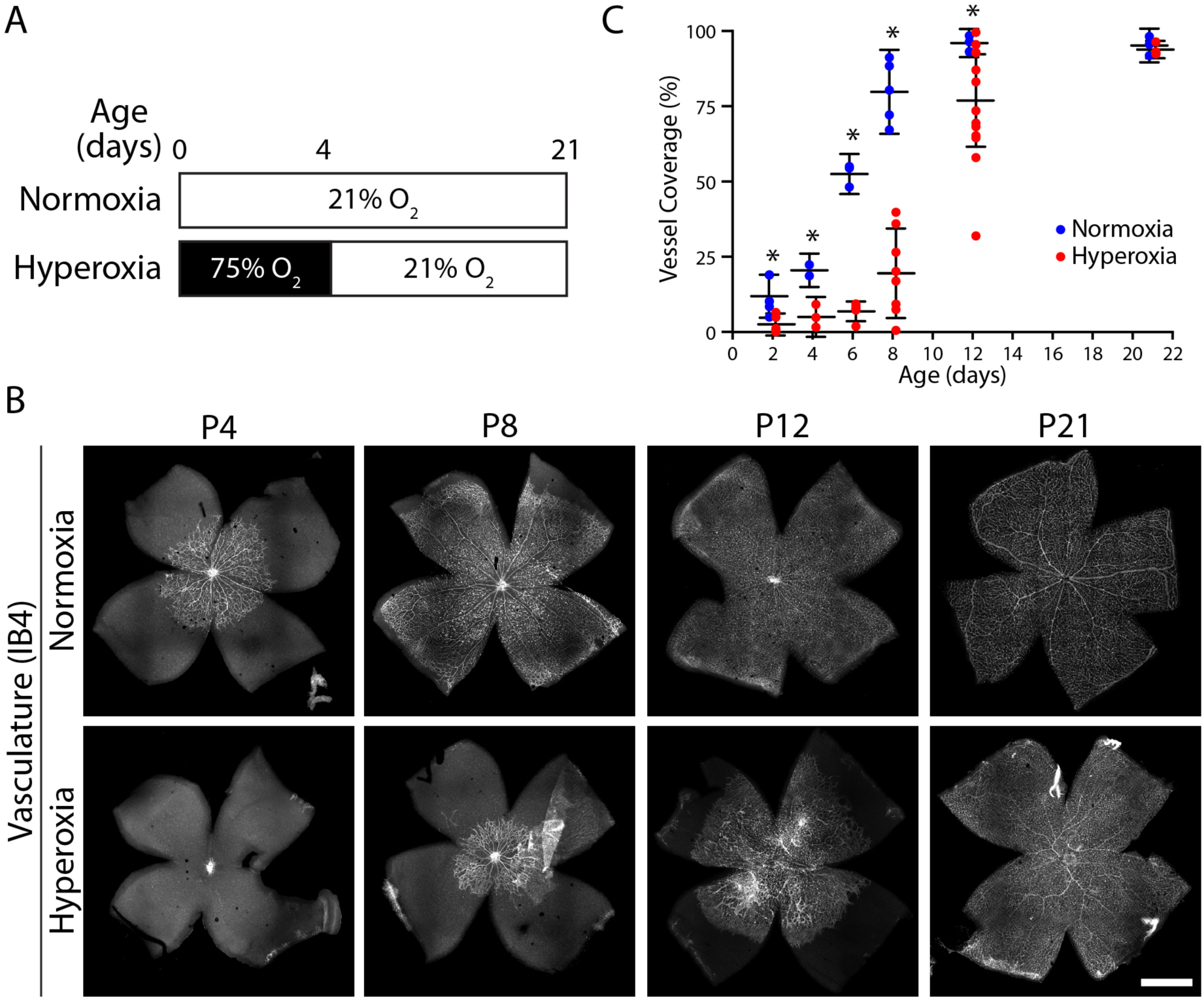
Neonatal hyperoxia prevents retinal angiogenesis. **A**) Schematic of experimental timeline. *Normoxia* (control) animals were maintained in room air (21% O_2_) throughout; *Hyperoxia* animals were raised in 75% O_2_ from P0-P4 before returning to room air (21% O_2_) at P4. **B**) Example confocal images of IB4-stained flat-mounted retinas, showing retinal vasculature. Images are stitched composites of individual 20x tiles. In normoxia controls, angiogenesis is initiated shortly after birth and the inner retinal surface is completely vascularized by P12. In the hyperoxia condition, angiogenesis is completely prevented until animals are returned to room air at P4. Scale bar: 1 mm. **C**) Summary data of retinal vascular area by condition for animals sacrificed at different ages. There was a significant angiogenic delay in hyperoxia animals. Statistics: Two-way ANOVA. Main effect of age F(5,50) = 82.6, p < 0.0001; main effect of oxygen F(1,50) = 55.7, p < 0.0001; interaction, F(5,50) = 9.4, p < 0.0001.). Asterisks denote significant differences by Holm-Sidak multiple comparisons test (p < 0.05). Error bars, mean ± s. d.

To test whether hyperoxia can suppress angiogenesis at later stages of its progression, we next exposed mice to 75% O_2_ starting at P2 or P4 and continuing for 2-4 days thereafter. At P2 and P4, retinal angiogenesis is underway but the vascular wavefront has not yet reached the periphery (Fig. 1B,C). In contrast to P0 exposure, we found that later O_2_ exposure did not delay the peripheral extension of the vascular wavefront, but instead resulted in central capillary loss reminiscent of the widely used mouse OIR paradigm (Supplemental Fig. S1). Together, these experiments show that early hyperoxia exposure can delay the onset of angiogenesis, but later exposure does not prevent the progression of angiogenic sprouting.

A close examination of the vascular phenotypes in P0-P4 hyperoxia-treated mice revealed that angiogenesis was not just delayed but also dysfunctional. We noted three particularly striking vascular abnormalities. First, about half of the eyes from P12 and older mice had conspicuous vitreous hemorrhages (Fig. 2A,B), which was never seen in controls. The presence of vitreous hemorrhage was also correlated with the severity of vascular delay at P12: eyes with bleeding showed greater delays in retinal vascular coverage (Fig. 2B). Second, the hyaloid system of hyperoxia treated eyes failed to regress. Dense hyaloid vessels were evident throughout the vitreous during dissection; the hyaloid artery could be seen emerging from the optic nerve head (Fig. 2C); and IB4-positive vascular tissue originating from the hyaloid vessels was present circumferentially in the peripheral retina (Fig. 2D). Third, all treated retinas, regardless of vessel coverage delay, exhibited abnormal vascular morphology: The radial organization of large vessels was usually disrupted; capillaries were of abnormally large caliber; and endothelial cells frequently formed lawns without any evident tubular structure (Fig. 2D,E). Moreover, vessel network organization was haphazard, with an apparent increase in vessel tortuosity and smaller capillary loops (Fig. 2D,E). Overall, these experiments demonstrate that neonatal hyperoxia not only delays angiogenesis but also causes vascular pathology in the developing mouse retina. Our results therefore establish the P0-P4 hyperoxia protocol as an experimental paradigm for neonatal oxygen induced retinopathy (NOIR).

**Figure 2.**
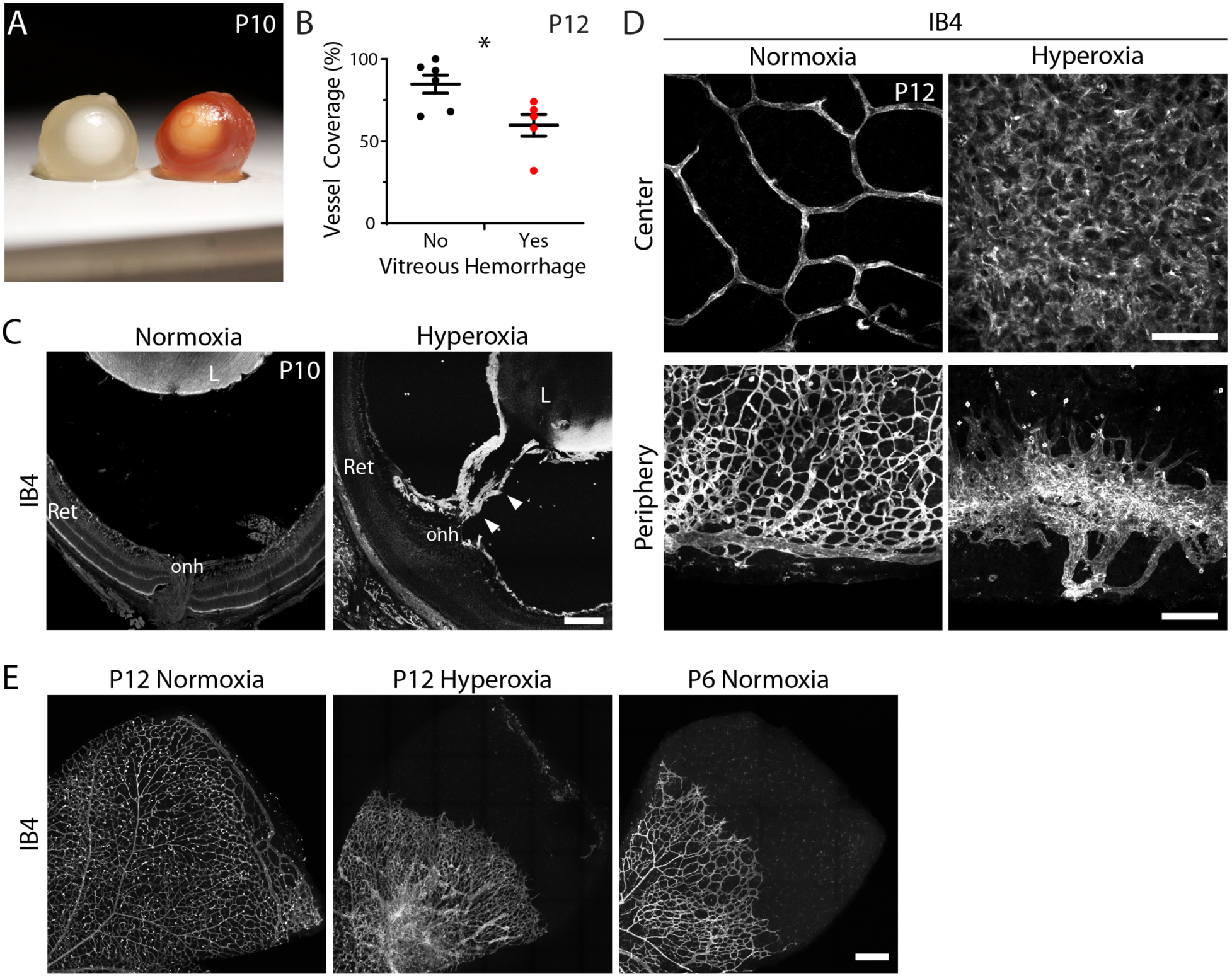
Angiogenesis is abnormal following return to room air in NOIR protocol. **A**) Photographs of freshly dissected P10 eyes from P0-P4 hyperoxia treated CD-1 mice, showing examples of globes with gross vitreous hemorrhage (right) and without hemorrhage (left). Note that CD-1 is an albino strain so hemorrhage is particularly obvious. **B**) Percentage of retinal area inside vessel wavefront in P12 NOIR retinas. Animals grouped by presence or absence of vitreous hemorrhage. **C**) Central retina region from P10 whole-eye cryosections stained with IB4 to reveal vasculature. Hyaloid vessels (arrows) traversing space between retina (Ret) and lens (L) are retained in hyperoxia but not control animals. ONH, optic nerve head. **D**) En-face confocal images of IB4-stained P12 retinas, chosen as representative examples of animals with macroscopic hemorrhage (see B). Images are Z-projections of stacks encompassing the nerve fiber layer. Controls (left) show mature capillary network in central retina, and complete primary plexus in the periphery with terminal circumferential vessel. Hyperoxia animals (right) show lawn of endothelial cells lacking capillary morphology in central retina, and periphery with IB4+ ring of vitreous-derived vascular tissue not connected to intrinsic retinal vessels. **E**) Low magnification en-face images illustrating vascular disorganization in hyperoxia treated mice. Images are oriented with ONH down and left. In P12 control, complete primary plexus is organized with large radial vessels giving rise to capillary network. P12 hyperoxia retina exhibits peripheral avascular zone, absence of large radial vessels, and hyperdense and irregular vascularity centrally. P6 control retina in which the vascular wavefront is at a similar eccentricity demonstrating that the P12 hyperoxia morphology does not resemble a more immature stage. Statistics: two-tailed T test (No hemorrhage, 84.7 ± 5.5% vascular coverage, n = 6; vitreous hemorrhage, 59.6 ± 6.6%, n = 5; *p = 0.026). Scale bars: C 100 µm; D top 50 μm; C bottom 150 μm; E 250 μm. Error bars, mean ± s. d.

### Hyperoxia suppresses and return to normoxia stimulates astrocyte proliferation

We next used the NOIR protocol to examine the effects of hyperoxia on developing astrocytes. To this end we stained retinal wholemounts with the astrocyte nuclear marker Sox9. The hyperoxic phase of the protocol – i.e. P0-P4 – corresponds to the period when astrocytes are normally migrating to colonize the retinal periphery (Chan-Ling et al., 2009; Fruttiger, 2002; O’Sullivan et al., 2017). This migration was unaffected by hyperoxia: astrocytes spread outward from the ONH to cover the retina, associated with RGC axons, and became polarized along the centrifugal axis similarly in normoxic and hyperoxic conditions. Accordingly, total astrocyte number was normal in treated mice at the conclusion of the hyperoxic phase (Fig. 3B). Thus, the aforementioned delay in angiogenesis (Fig. 2) was not due to impairment in astrocyte colonization of the retina during the P0-P4 period.

**Figure 3.**
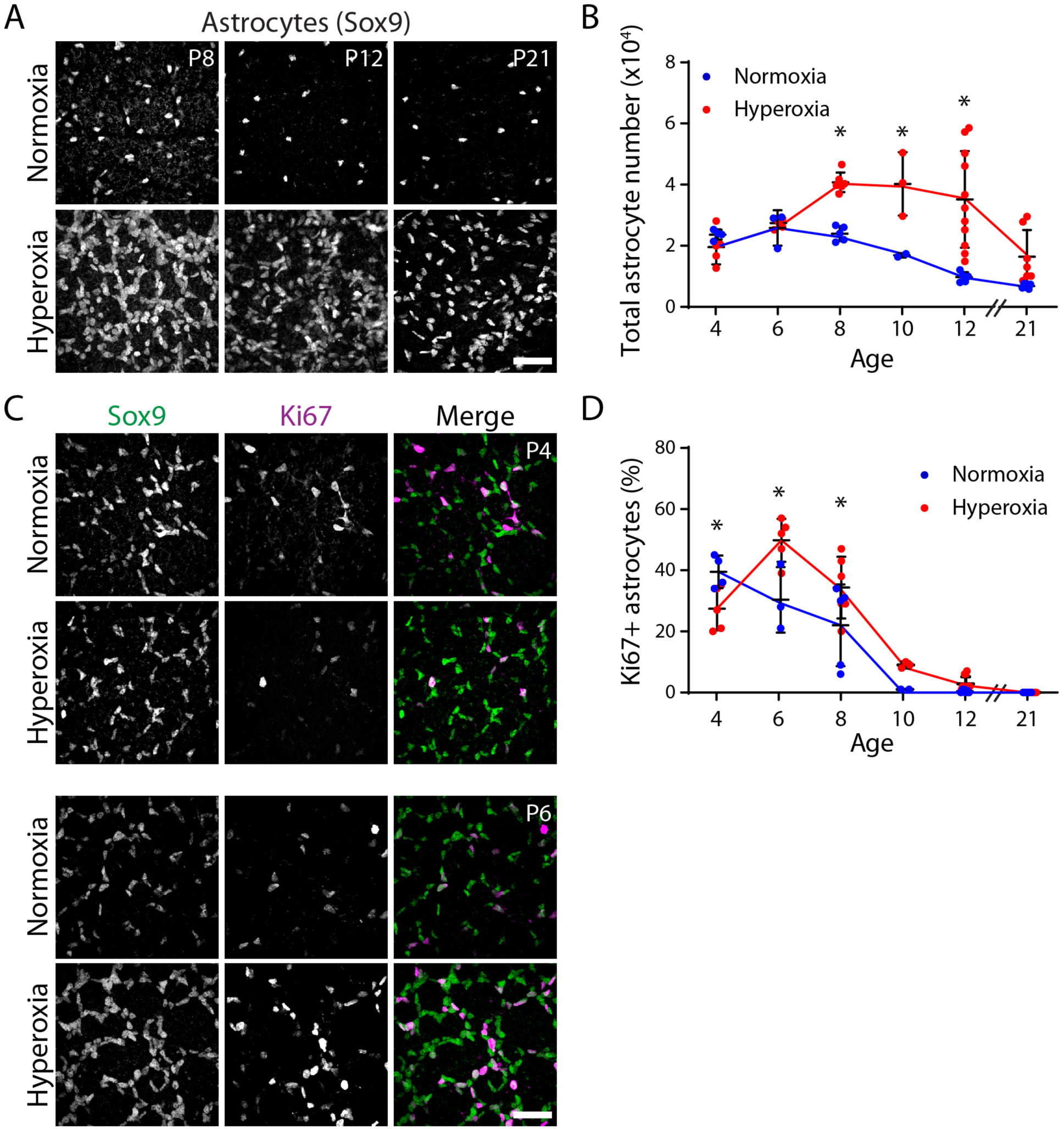
Astrocyte proliferation is suppressed by hyperoxia and stimulated by return to room air. **A**) Example en-face confocal images of the astrocyte nuclear marker Sox9. Hyperoxia treated retinas show significantly denser astrocytes. **B**) Summary of total astrocyte numbers calculated from quantification of images such as those shown in (A). From P8 to P12, hyperoxia exposed retinas had significantly more astrocytes than controls. At P21 there was not a group difference but some individual hyperoxia animals have nearly triple the number of astrocytes as controls. **C**) Example en-face images of Sox9 and Ki67 double-stained retina. Ki67 marks mitotically active cells. Double-positive cells were counted as dividing astrocytes. **D**) Quantification of astrocyte proliferation. At P4, the percentage of mitotically active astrocytes was reduced in hyperoxia retinas compared to controls, whereas at P6,-8 after return to normoxia, more astrocytes were actively dividing compared to controls. Statistics: two-way ANOVA. B: Main effect of age F(5,46) = 11.67, p < 0.0001; main effect of oxygen F(1,46) = 29.85, p < 0.0001; interaction, F(5,46) = 4.999, p = 0.001. D: Main effect of age F(5,49) = 74.5, p < 0.0001; main effect of oxygen F(1,49) = 7.9, p = 0.007; interaction, F(5,49) = 6.336, p = 0.0001. Asterisks denote significant differences by Holm-Sidak post-hoc test. P-values in B: P8, p = 0.082; P10, p = 0.0136. P12, p < 0.0001. P-values in D: P4, p = 0.0268; P6, p = 0.0007; P8, p = 0.0122. Scale bars 50 μm. Error bars, mean ± s. d.

While the astrocyte template was largely unaffected by hyperoxia, we found that the hypoxic stress associated with the return to room air significantly perturbed astrocyte development. Whole-mount staining for astrocyte nuclear markers revealed that, four days after return to room air, treated mice exhibited a striking increase in astrocyte numbers (Fig. 3A). In wild-type mice, astrocyte numbers normally increase until P5-6 due to ongoing migration and proliferation; subsequently, their numbers decline substantially due to cell death (Bucher et al., 2013; Chan-Ling et al., 2009; Puñal et al., 2019). A similar pattern was observed in our control normoxia mice (Fig. 3B). By contrast, in treated mice the number of astrocytes began to increase after P6, during the time when the astrocyte population was dwindling in controls (Fig. 3A,B). At P12, treated animals had on average 3.6-fold more astrocytes than normoxic controls, with some animals showing even more dramatic effects (Fig. 3B). Astrocyte numbers remained elevated for weeks – in some cases until at least P21 (Fig. 3B).

Given these large differences in astrocyte numbers, we surmised that astrocyte mitotic activity might be regulated by oxygen. To test this idea, we used Ki67 as an immunohistochemical marker of proliferating cells (Fig. 3C). In normoxic controls, the fraction of Sox9^+^Ki67^+^ proliferative astrocytes decreased monotonically over development, such that virtually all astrocytes were quiescent by P10 (Fig. 3D). By contrast, in mice exposed to the NOIR protocol, astrocyte proliferation was regulated in a triphasic manner: Initially, during the hyperoxic phase, astrocyte proliferation was suppressed relative to controls; the return to room air then stimulated increased rates of proliferation at P6 and P8 before mitotic activity ultimately fell to control levels (Fig. 3C,D). Importantly, the increase in proliferative astrocytes at P6 preceded the increase in astrocyte number at P8, suggesting that proliferation likely accounts for the expansion of the astrocyte population. These findings strongly suggest that return to room air at P4 stimulates astrocyte proliferation, thereby elevating astrocyte numbers in NOIR mice. The proliferative response appears to be reliable across mice, as both the surge in proliferation at P6 (Fig. 3D) and the surge in astrocyte abundance at P8 (Fig. 3B) were highly consistent between animals. Subsequent factors affecting astrocyte number, by contrast, varied substantially between individual mice given the wide range in astrocyte numbers that emerges by P12 (Fig. 3B,D).

### Astrocyte patterning following hyperoxia is defective

The observation that astrocyte numbers are abnormal in NOIR mice raised the possibility that the angiogenic astrocyte template is disturbed by early hyperoxia. To investigate how the NOIR protocol affects astrocyte patterning, we evaluated soma positioning and arbor anatomy in retinal whole-mounts, using antibody markers that revealed either the nucleus (Sox9) or the arbors (GFAP, PDGFRα) of developing astrocytes. This analysis revealed that, in P12 treated retinas, astrocytes are arranged into irregular clumps and strands instead of being evenly spaced (Fig. 4A). The most striking effect was ahead of the vascular wavefront, where chains of astrocyte somata were typically arranged into large polygons that left large swaths of retinal territory uncovered by astrocytes or their arbors (Fig. 4A,B). Because the pattern of angiogenesis normally follows the astrocyte template, dysmorphia in the astrocyte network may be transmitted to growing vessels. Consistent with this possibility, endothelial tip cells and their filopodia at the vascular wavefront remained strictly co-localized with astrocytes in NOIR mice, even when astrocyte distribution was disturbed (Fig. 4B). Together, these findings suggest that oxygen stress alters the astrocyte template in a manner that could result in delayed and/or disrupted angiogenesis.

**Figure 4.**
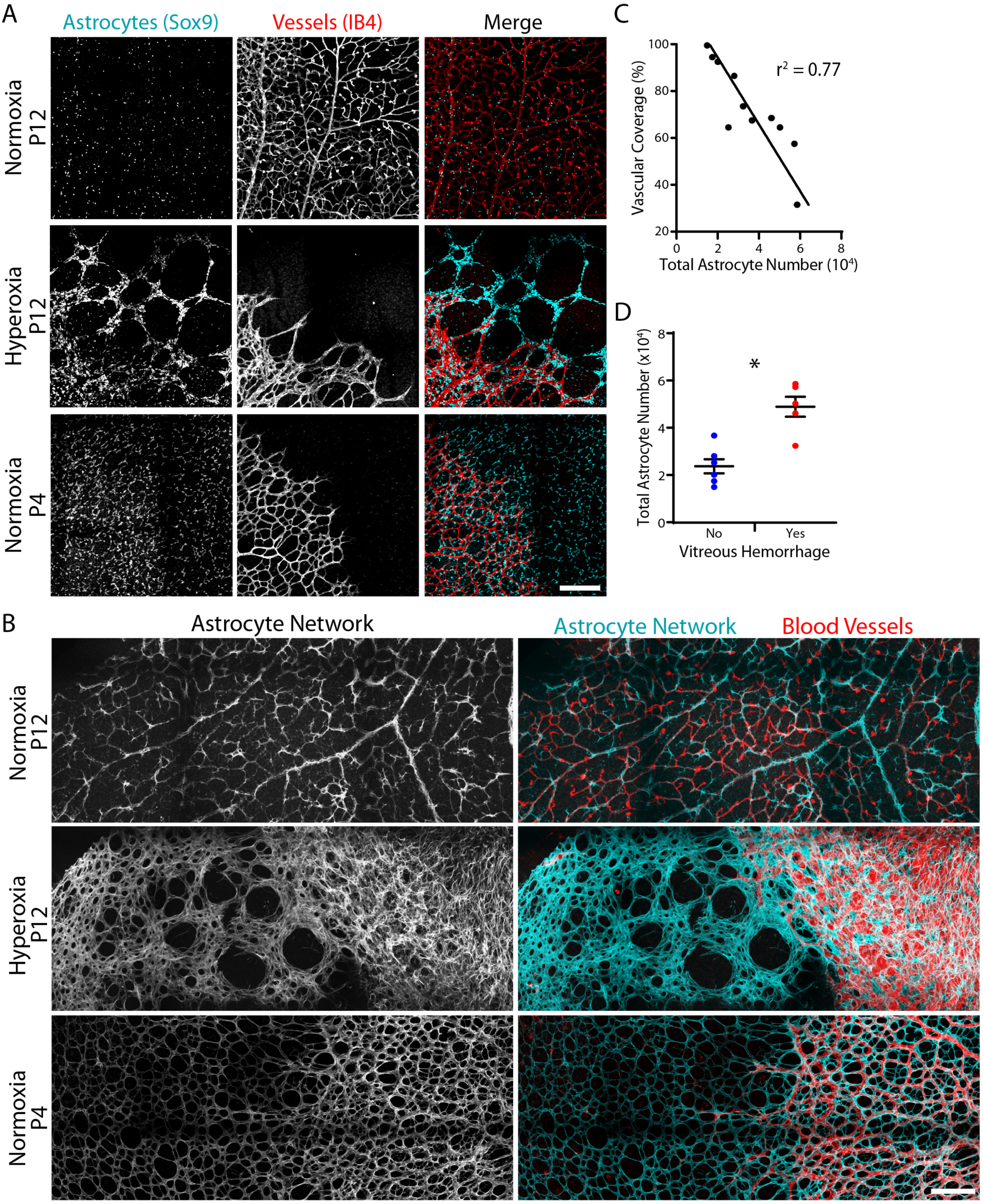
Astrocyte patterning is abnormal after hyperoxia. **A**) Representative en-face confocal images of midperipheral retina stained with Sox9 and IB4 to label astrocyte nuclei and vasculature. **B**) Representative en-face images of astrocyte arbors, labeled by anti-GFAP (P12) or anti-PDGFRα (P4). In P12 controls, astrocytes are homogenously distributed and evenly spaced across the retina (A,B). In P12 hyperoxia retinas, astrocyte somata are aggregated in clumps and strings in advance of the delayed vascular wavefront (A). Uneven patterning of the astrocyte arbor network is also evident (B). Despite these patterning defects, advancing endothelial cells remain associated with astrocytes, and do not enter astrocyte-free retinal regions (A,B). Blood vessels have reached a similar eccentricity in P4 controls, but astrocytes are evenly distributed rather than displaying the aggregation seen in hyperoxia retinas (A). The P4 arbor network is also far more uniform than in P12 treated animals (B). **C**) In P12 hyperoxia treated retinas, retinal vascular coverage is inversely correlated with the total number of astrocytes. **D**) Hyperoxia retinas with vitreous hemorrhages (e.g. Fig. 2A) have significantly more astrocytes than hyperoxia retinas without hemorrhage. Statistics, two-tailed T test; no hemorrhage, 23,721 ± 2,982 total astrocytes, n = 6; vitreous hemorrhage, 48,895 ± 4,220, n = 5; p = 0.0015. Scale bars 200 μm. Error bars, mean ± s. d.

### Astrocyte number predicts vascular abnormalities

The finding that neonatal hyperoxia causes both vascular and glial abnormalities spurred us to investigate whether the vascular defects might in fact originate with the glia. To this end we exploited the variability in the number of astrocytes in hyperoxia-exposed retinas at P12 (Fig. 3B). If vessel phenotypes are a consequence of astrocyte phenotypes, we would predict that the NOIR animals with the most severe astrocyte disruptions should also show the most severe vascular pathology. Consistent with this model, the total number of astrocytes was inversely correlated with vascularized retinal area, such that retinas with the greatest expansion of the astrocyte population displayed the most profound delay in peripheral extension of retinal vessels (Fig. 4C). Furthermore, animals with vitreous hemorrhage had significantly more astrocytes than those that escaped this pathology (Fig. 4D). These observations support the notion that anomalous astrocyte proliferation promotes angiogenic defects in the NOIR paradigm.

### Neonatal hyperoxia causes enduring retinal abnormalities

To explore the long-term consequences of exposure to neonatal hyperoxia, we examined NOIR mice and normoxic controls at three weeks of age, when the retina is largely mature. Vascular disorganization and persistent hyaloid vessels were still evident in whole-mount NOIR retinas at P21, particularly in cases where astrocyte numbers remained high (Fig. 5C). To test for other facets of retinal pathology, retinas were cryosectioned and immunostained with vascular and glial markers to assess their cytoarchitectural integrity. Hyperoxic retinas (n = 4) showed a strikingly abnormal cross-sectional profile, characterized by buckling and corrugation of the outer layers (Fig. 5A) in a manner reminiscent of pathology seen in some samples from eyes with advanced ROP (Foos, 1987). This was accompanied by large-scale reactive gliosis of Müller glia, indicated by upregulation of GFAP (Fig. 5B), which is consistent with the presence of ongoing cellular insults or tissue stress. We also noted additional vascular phenotypes in IB4-stained sections that were not evident in the whole-mount preparations. These included laminar disorganization of the intermediate and deep vascular plexuses (Fig. 5A); intravitreal neovascularization (Fig 5B); and hyperplasic vascular tissue at the inner retinal surface (Fig. 5A). The hyperplasic superficial vasculature was associated with a hyperplasic GFAP+ glial network, which likely included excess nerve fiber layer astrocytes as well as the endfeet of reactive Müller glia (Fig. 5B). Together, these observations indicate that a brief period of neonatal hyperoxia during a crucial period of glial-vascular development produces long-lasting deleterious consequences.

**Figure 5.**
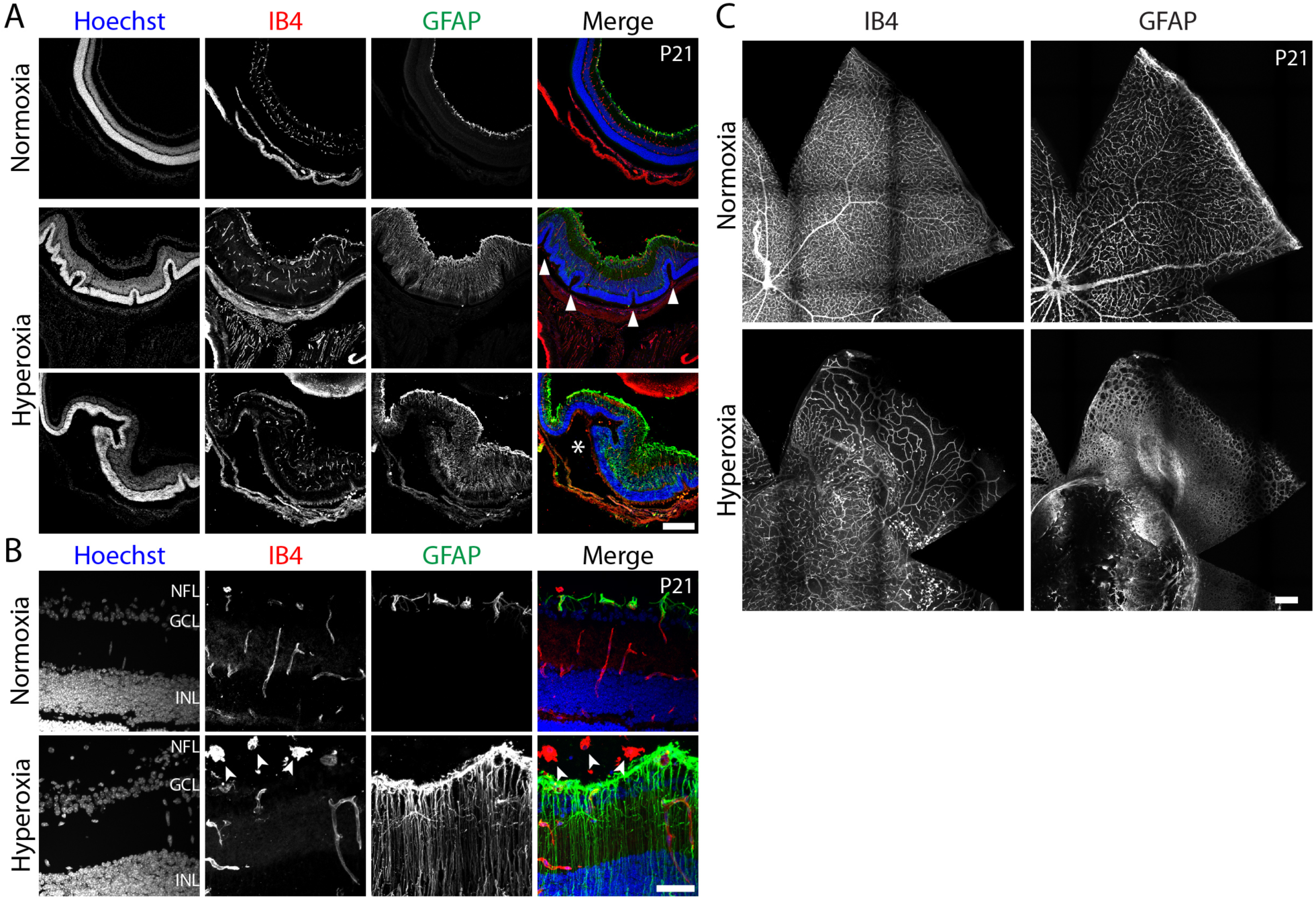
Neonatal hyperoxia causes enduring retinopathy. **A**,**B**) Retinal cross-sections from P21 mice illustrating pathological features caused by P0-P4 hyperoxia. Low (A) and high (B) magnification representative confocal images of control or hyperoxia tissue stained for vasculature (IB4) and GFAP. Two examples of different hyperoxia retinas are given in A. Controls display regular laminar organization with smooth contours. GFAP staining is restricted to inner retina where astrocytes reside in the RNFL. Hyperoxia retinas are thickened; show ROP-like outer retinal folding (arrowheads; Foos, 1987); and express GFAP throughout the full depth of the retina indicating reactive Müller gliosis. Asterisk indicates large subretinal space underlying folded neuroretina suggestive of retinal detachment. B) Higher magnification view shows vitreal IB4^+^ neovascular clumps, and radial distribution of GFAP staining in hyperoxia retinas consistent with Müller glia labeling. **C**) En-face view of vascular (IB4) and glial (GFAP) phenotypes at P21. Scale bars: A,C) 250 μm; B) 50 μm.

### Hypoxia stimulates astrocyte proliferation

Because neonatal hyperoxia causes long-lasting astrocyte and vascular pathology in the retina, we decided to investigate what mechanism could produce such effects. Our initial results suggested a central pathogenic role for aberrant astrocyte proliferation (Fig. 3), so we focused on identifying the mechanisms that stimulate astrocyte overproduction in NOIR mice. It was clear from the NOIR experiments that exposure to high oxygen suppresses astrocyte proliferation, while return to room air promotes proliferation (Fig. 3D). These observations led us to hypothesize that the state of the cellular hypoxia-sensing machinery might directly regulate astrocyte proliferation. In this model, the cause of proliferation in the NOIR protocol is the relative hypoxia experienced by the retinal tissue upon transition from 75% to 21% oxygen. If this hypothesis is correct, then rearing mice in a hypoxic environment should mimic the proliferative effects of return to room air in the NOIR protocol. To test this idea, neonatal mice were raised in 10% O_2_ from P0-4. At both P2 and P4, hypoxic mice showed a striking increase in the fraction of Ki67^+^ proliferating astrocytes (Fig 6A,B). This was accompanied by a corresponding increase in astrocyte abundance: By P4, astrocyte density in hypoxic retinas exceeded controls by approximately two-fold (Fig. 6C). Therefore, direct hypoxia drives astrocytes to divide in a manner similar to the room-air phase of the NOIR paradigm, suggesting that a relative decrement in oxygen availability is the physiologic stimulus that drives astrocyte mitosis.

**Figure 6.**
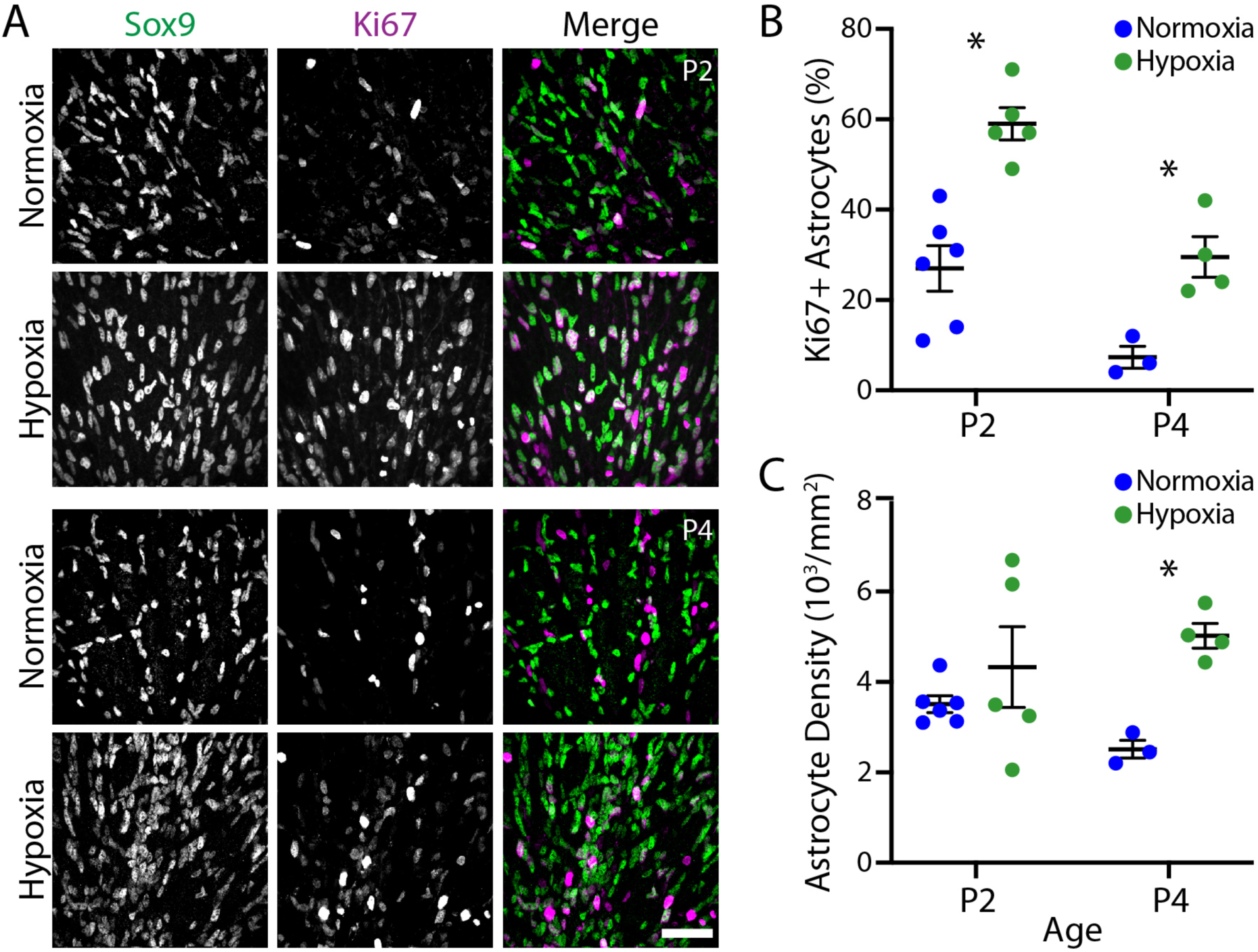
Hypoxia stimulates astrocyte proliferation. **A**) En-face confocal images showing co-labeling for Sox9 and Ki67 in animals exposed to 10% hypoxia from birth, or their littermate controls. Representative examples are shown at P2 and P4. **B**) Effects of hypoxia on astrocyte proliferation were quantified from images similar to A, by calculating the fraction of astrocytes that expressed Ki67. Astrocyte proliferation was higher at both P2 and P4 in treated mice. Statistics: Two-way ANOVA. Main effect of age F(1,14) = 27.49, p = 0.0001; main effect of oxygen F(1,14) = 33.37, p < 0.0001 ; interaction, F(1,14) = 1.10, p = 0.3121. **C**) Effects of hypoxia on astrocyte abundance, calculated from images similar to A. Astrocyte density became higher by P4 in treated animals. Statistics: Two-way ANOVA. Main effect of age F(1,14) = 0.07, p = 0.7885; main effect of oxygen F(1,14) = 9.00, p = 0.0095 ; interaction, F(1,14) = 2.32, p = 0.1502. Asterisks denote significant differences by post-hoc Holm-Sidak test. B: P2, p = 0.0002; P4, p = 0.0184. C: P2, p = 0.4434; P4, p = 0.0238. Scale bar, 50 µm. Error bars, mean ± s. d.

### Astrocyte HIF2α drives proliferation in normal development and following hyperoxia

Finally, we investigated the molecular and cellular mechanisms that promote astrocyte proliferation upon exposure to relative hypoxia. The hypoxia-inducible factor (HIF) pathway is a key molecular sensor of tissue oxygen levels that contributes to angiogenesis, both in brain (Zhang et al., 2020) and retina (Duan et al., 2014; Kurihara et al., 2010; Nakamura-Ishizu et al., 2012; Weidemann et al., 2010). Developing retinal astrocytes strongly express *Epas1*, which encodes the HIF2α transcription factor, indicating that astrocytes possess the molecular machinery to detect tissue hypoxia (B. Clark, personal communication; Clark et al., 2019). Furthermore, HIF signaling is clearly active within astrocytes during retinal development because astrocytes express HIF target genes – such as *Vegfa* – in a hypoxia-dependent manner (Gerhardt et al., 2003; Pagès and Pouysségur, 2005; Stone et al., 1995; West et al., 2005). We therefore hypothesized that HIF-mediated oxygen sensing occurs within astrocytes to promote their proliferation in a cell-autonomous fashion. To test this idea we used a conditional knockout strategy, in which mice carrying a floxed HIF2α allele (Gruber et al., 2007) were crossed to astrocyte-specific *GFAP-Cre* mice (O’Sullivan et al., 2017; Puñal et al., 2019; Zhuo et al., 2001). This breeding yielded <u>a</u>strocyte-specific <u>c</u>onditional <u>HIF2α k</u>nock<u>o</u>ut animals (abbreviated AC-Hif2α-KO).

Using these mutants, we first investigated the effects of astrocyte HIF2α deletion under normoxic conditions. Although previous studies have addressed this question, the precise role of HIF2α in astrocytes has remained controversial (Duan et al., 2014; Weidemann et al., 2010). We found that removal of HIF2α was sufficient to abrogate HIF-dependent gene expression, as shown by immunostaining for the HIF target VEGF-A (Fig. 7A,B). In wild-type retinas at P2, the vast majority of astrocytes (92.1%) expressed VEGF-A (n = 2151 cells from 3 mice). While expression levels differed between vascularized central and avascular peripheral regions (Fig. 7A), a similar fraction of astrocytes were VEGF^+^ regardless of retinal location (88.7% in central retina, 96.0% middle retina, 90.1% peripheral retina). By contrast, in P2 AC-Hif2α-KO mice, most retinal astrocytes lacked VEGF-A immunoreactivity (Fig. 7B), suggesting that HIF2α is the major effector of HIF signaling in retinal astrocytes. A subset of AC-Hif2α-KO astrocytes remained VEGF-A^+^ (Fig. 7B), consistent with our previous observations that a small fraction of cells escape Cre recombination in *GFAP-Cre* mice (O’Sullivan et al., 2017; Puñal et al., 2019). Together, these findings indicate that removal of HIF2α blocks HIF pathway activity in a cell-autonomous manner.

**Figure 7.**
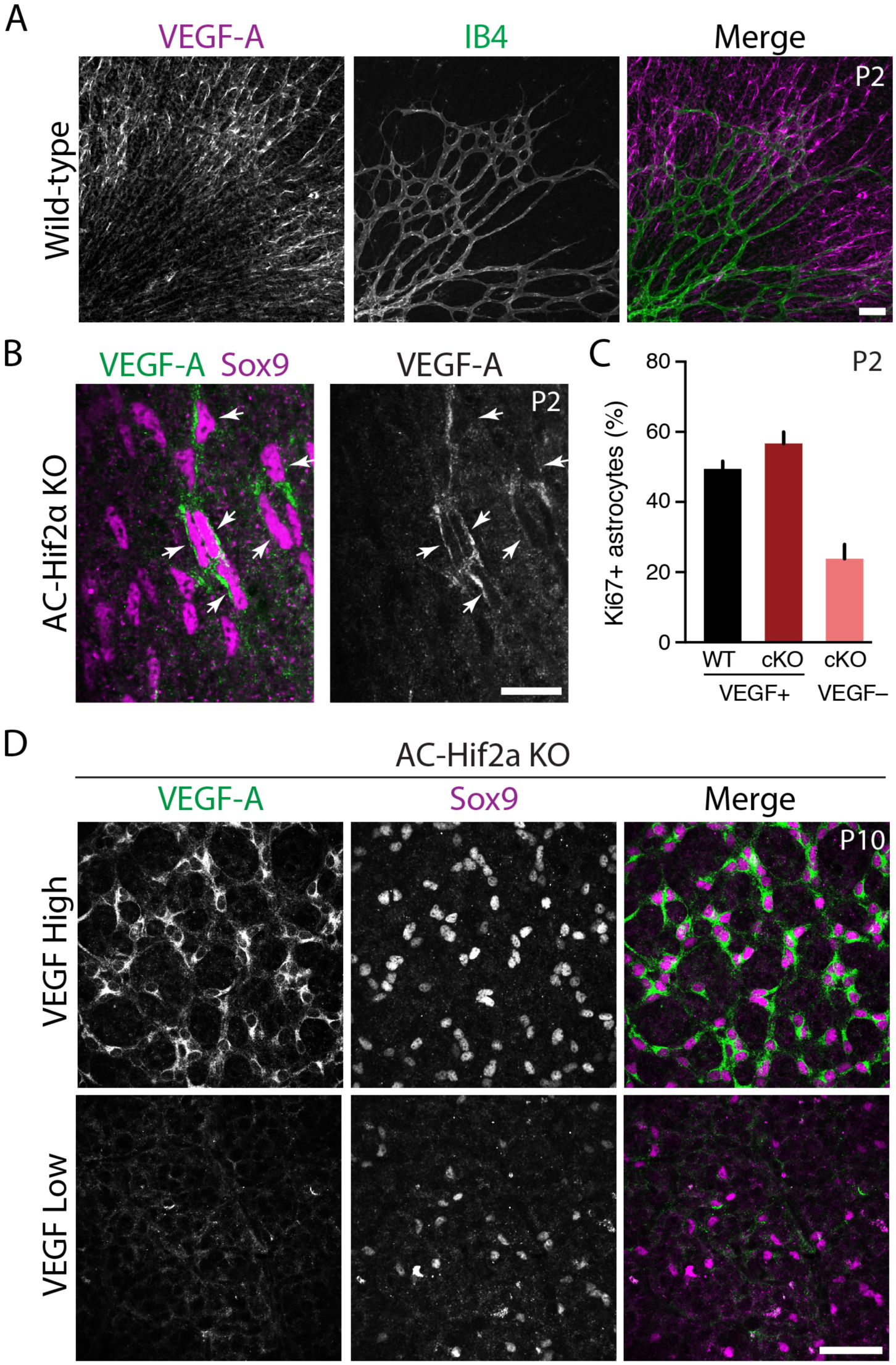
Astrocyte-specific deletion of Hif2α blocks HIF signaling and impairs astrocyte proliferation. **A**) P2 wild-type retinal whole-mounts stained for VEGF-A and vasculature (IB4). Retinal astrocytes in avascular peripheral regions have higher VEGF-A expression, but astrocytes still express VEGF-A within vascularized central retina. This expression pattern is consistent with previous studies that used in situ hybridization as a *Vegfa* readout (Gerhardt et al., 2003; West et al., 2005). **B**) In P2 *GFAP-Cre;* Hif2α^*flox/flox*^ mice (AC-Hif2α-KO), many astrocytes fail to express VEGF-A, indicating loss of HIF signaling. A subset of astrocytes remains VEGF-A^+^ (arrowheads). **C**) Quantification of astrocyte proliferation in wild-type (WT) and mutant retinas triple stained for Sox9, VEGF-A, and Ki67. In AC-Hif2α-KO mutants, Sox9^+^ astrocytes without HIF signaling (VEGF^−^) are significantly less proliferative than those retaining HIF signaling (VEGF^+^). Mutant VEGF^+^ astrocytes proliferate at a similar rate as WT astrocytes. Error bars: 95% confidence interval. N = 3 animals per genotype. **D**) Two classes of mutant phenotypes in P10 AC-Hif2α-KO mice. VEGF-high mutants express VEGF-A in all astrocytes (top); in VEGF-low mutants (bottom), VEGF-A is virtually absent from astrocytes. Note also the difference in astrocyte cell density between the two mutant classes (also see Fig. 8C and Supplemental Fig. S2 for quantification of astrocyte density in the different mutant classes and littermate controls).

To learn whether astrocyte proliferation is affected by loss of HIF signaling, we evaluated the number of Ki67^+^ proliferating astrocytes at P2 in AC-Hif2α-KO mutants and littermate controls. In mutants, we separately quantified the two distinct astrocyte populations – i.e. VEGF-A^−^ astrocytes that had lost HIF signaling, and VEGF-A^+^ astrocytes in which HIF signaling was intact (Fig. 7B). This analysis revealed a significant proliferation deficit in HIF-deficient astrocytes relative to VEGF-A^+^ internal controls, or to wild-type astrocytes from littermates (Fig. 7C). The decreased proliferation rate in mutants was accompanied by a large decrement in mutant astrocyte numbers at P2 (astrocyte number per control retina: 13,068 ± 1,705, n = 3; astrocyte number per mutant retina: 7,548 ± 2,070, n = 4; mean ± s.d.). These findings are consistent with a previous study that showed a severe reduction in astrocyte numbers upon astrocyte-specific Hif2α deletion (Duan et al., 2014). Overall, these data are all consistent with the notion that Hif2α is required for normal astrocyte proliferation.

Given these straightforward results, we were surprised to find that the AC-Hif2α-KO phenotype became more complex at later developmental stages. At P10, staining for VEGF-A and for astrocyte markers revealed two distinct classes of mutants (Fig. 7D; Supplemental Fig. S2). The first class of mutants replicated the findings of the Duan et al. (2014) study: Astrocyte numbers were reduced (Fig. 7D; Supplemental Fig. S2) and retinal vasculature was entirely absent. In this class of mutants, which we denote “VEGF-low”, astrocyte VEGF-A staining was almost entirely absent, demonstrating effective elimination of HIF2α function (Fig. 7D). By contrast, in the second class of mutants denoted “VEGF-high,” virtually all astrocytes continued to express VEGF-A (Fig. 7D). This was surprising because, in P2 mutants, we always observed a mixture of VEGF-A^+^ and VEGF-A^−^ astrocytes (Fig. 7B). In VEGF-high animals, retinal vasculature was present and relatively normal, albeit occasionally delayed in its development, and astrocyte numbers were similar to wild-type (Supplemental Fig. S2 and data not shown). The VEGF-high phenotype is consistent with the proliferative defect observed at P2: Because VEGF-A^+^ (presumably Cre-negative) astrocytes have a proliferative advantage over their mutant neighbors (Fig. 7C), these wild-type cells may in some cases outcompete the mutant astrocytes to take over the entire retina by P10. Therefore, the phenotypes observed in P2 and P10 AC-Hif2α-KO mutants together support the conclusion that HIF signaling is required to support astrocyte proliferation during normal retinal development.

Next we tested whether astrocyte HIF function mediates pathological proliferation upon exposure to relative hypoxia. To address this question we subjected AC-Hif2α-KO mice and their littermate controls to the NOIR paradigm. Littermates included both HIF2α^*flox/flox*^ mice lacking the Cre transgene and *GFAP-Cre;* HIF2α^*+/+*^ mice, which were phenotypically indistinguishable; together we refer to them as HIF2α^*WT*^ animals. AC-Hif2α-KO mice exhibiting the VEGF-high phenotype (as described above) were excluded from the analysis; it was clear from the VEGF staining pattern that this class of mutants did not lack astrocytic HIF2α function (Fig. 8A). As in our previous experiments using CD-1 mice (Fig. 3), HIF2α^*WT*^ mice exposed to the NOIR protocol showed a dramatic increase in astrocyte numbers at P10. By contrast, no such increase was observed in AC-Hif2α-KO mice (Fig. 8B,C; Supplemental Fig. S2B). These data strongly suggest that HIF2α is required within astrocytes to drive the proliferative response observed following return to room air at P4.

**Figure 8.**
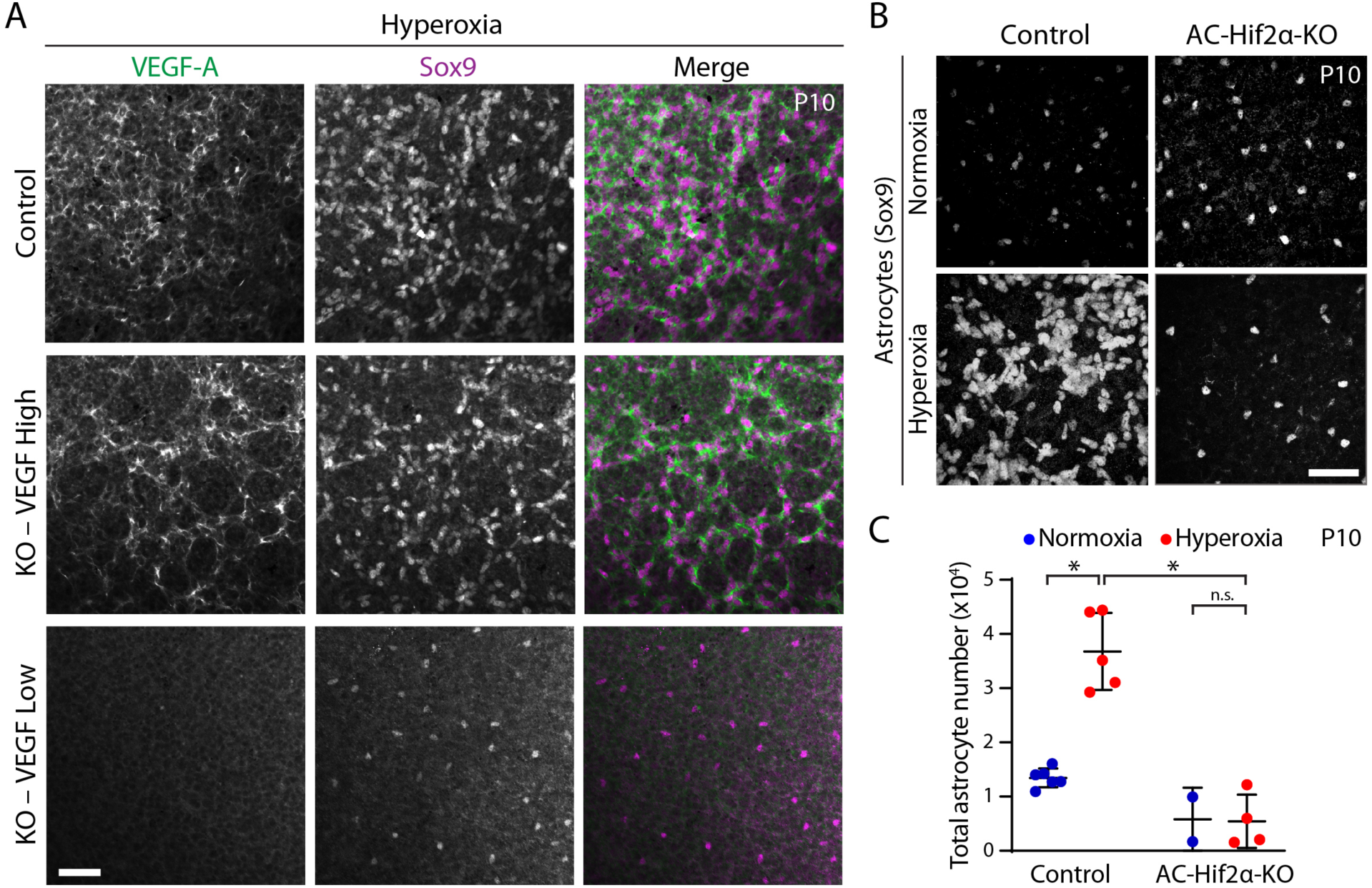
Astrocyte HIF2α is required for proliferation on return to room air in NOIR paradigm. **A**) Characterization of HIF pathway activation in astrocyte-specific *GFAP-Cre;* Hif2α^*flox*^ mutants (AC-Hif2α-KO). Representative en-face views of P10 retinas stained for astrocyte marker Sox9 and for anti-VEGF-A, a marker of HIF pathway activation. HIF2α^*WT*^ littermate controls (top row) show substantial astrocytic expression of VEGF in the NOIR paradigm. VEGF immunoreactivity is largely localized to Sox9^+^ cells and their arbors. In AC-Hif2α-KO mutants, we observed two phenotypic classes denoted VEGF High and VEGF Low. The first class (center row) exhibited VEGF staining indistinguishable from controls, suggesting a failure of HIF2α gene deletion. These were excluded from further analysis. VEGF Low mutants (bottom row) entirely lacked VEGF expression indicating successful abrogation of HIF signaling. **B**,**C**) Effects of the NOIR paradigm on P10 astrocyte numbers in AC-Hif2α-KO mice. B, representative Sox9 images; C, quantification of total Sox9^+^ astrocytes. Astrocyte numbers were greatly increased following hyperoxia in littermate controls, but not in AC-Hif2α-KO mutants. Statistics: Two-way ANOVA; main effect of O_2_ treatment F(1,13) = 19.0, p = 0.0008; main effect of genotype F(1,13) = 54.7, p = 5.2 × 10^−6^; interaction, F(1,13) = 20.2, p = 0.0006. Asterisks denote significant differences by post-hoc Holm-Sidak test. WT normoxia vs. hyperoxia, p = 1.7 × 10^−5^. WT hyperoxia vs KO hyperoxia, p = 2.3 × 10^−6^; KO normoxia vs. KO hyperoxia, p = 0.9354. Scale bars, 50 µm. Error bars, mean ± s. d.

## Discussion

In this study, we found that oxygen stress perturbs development of retinal astrocytes. Return to room air following a limited period of neonatal hyperoxia stimulated exuberant astrocyte mitotic activity, at an age when maturing astrocytes are normally ceasing their proliferation. Exposure to neonatal oxygen also perturbed retinal angiogenesis: We observed delays in peripheral vessel extension; persistent hyaloid vasculature; irregular and abnormally dense endothelial networks lacking regular capillary morphology; and associated vitreous hemorrhage. Many of these vascular phenotypes have features resembling ROP or other vitreoretinopathies (Eller et al., 1987; Foos, 1987; McMenamin et al., 2016). Within retinas, the severity of vascular pathologies was correlated with the total number of astrocytes, suggesting that the two phenotypes may be linked. We propose a model in which neonatal hyperoxia recalibrates the baseline oxygen level sensed by astrocytes, such that return to room air triggers a HIF2α-dependent hypoxic response that includes astrocyte proliferation. Subsequently, excessive astrocyte numbers contribute to developmental retinal vasculopathies including delayed peripheral extension, abnormal vascular patterning, and hemorrhage. These vascular pathologies are accompanied by pathological changes to overall retinal cytoarchitecture. Altogether, our findings raise the possibility that astrocytes have an important role in the pathobiology of ROP and related disorders.

### Retinal astrocyte proliferation is bidirectionally regulated by environmental oxygen

It has long been suspected that retinal astrocytes might be sensitive to tissue hypoxia, as astrocytes are the major source of VEGF-A in the neonatal retina and have a key role in promoting angiogenesis (Rattner et al., 2019; Stone et al., 1995). Furthermore, the arrival of blood vessels at any given retinal location induces key features astrocyte maturation: VEGF expression is downregulated while GFAP and other mature markers become upregulated (Chan-Ling et al., 2009; Duan et al., 2017; Gerhardt et al., 2003; West et al., 2005). These observations suggest that immature astrocytes have the capacity to detect tissue oxygen levels and are a key part of the feedback loop that drives angiogenesis in response to hypoxia. However, it was unclear whether this capacity for oxygen sensing is used as a mechanism to control astrocyte development. Previous studies addressing this question found only minor effects on astrocyte morphology or maturation when rodents were exposed to hyperoxia (Duan et al., 2017; Zhang et al., 1999). Here we show that mitotic activity of neonatal astrocytes is bidirectionally regulated by oxygen – hyperoxia from P0-P4 suppresses proliferation while direct hypoxia promotes proliferation. Our data are consistent with a model in which astrocytes sense local tissue oxygen levels using the HIF2α pathway, which drives proliferation in a manner that is proportional to the amount of HIF signaling. Such a mechanism would serve retinal metabolic needs by ensuring the addition of more VEGF-A^+^ angiogenic astrocytes when the tissue is hypoxic, and by suppressing this process once vessels have arrived to relieve local hypoxia. Manipulations of environmental oxygen appear to disrupt this homeostatic mechanism, such that decrements from baseline oxygenation are sufficient to drive HIF activity and excessive astrocyte proliferation. Consistent with this idea, we observed that both absolute hypoxia (10% O_2_) and normoxia following hyperoxia (dropping from 75% to 21% O_2_) can stimulate astrocyte proliferation in a similar manner.

West et al. (2005) reported a seemingly contradictory result, in that astrocytes in hyperoxia-exposed retinas had higher rates of BrdU incorporation compared to normoxic controls. In this study as in ours, hyperoxia prevented formation of retinal vasculature (Fig. 1). However, West et al. (2005) used a much longer 8-day hyperoxia exposure. Therefore, retinal neurons were deprived of an intrinsic blood supply for four days longer than in our NOIR paradigm. We suggest that 8 days of avascularity likely caused tissue hypoxia, with consequent astrocyte proliferation (Fig. 3), even while environmental oxygen was high. Indeed, the interpretation proposed by West et al. (2005) was that hypoxia – not hyperoxia – was driving the proliferation they observed.

It is likely that the pro-proliferative effect of hypoxia is limited to a brief period of astrocyte development, since no such proliferation has been reported following return to room air in the P7-12 OIR paradigm. By P7 retinal astrocytes are substantially more mature: They have completed migration, ceased proliferating, and assumed a mature GFAP-expressing molecular profile (Chan-Ling et al., 2009; O’Sullivan et al., 2017; West et al., 2005). It is plausible that mature astrocytes may respond differently to metabolic stresses. Furthermore, most retinal astrocytes reside in close contact with vasculature by P7 – another key change that could contribute to the difference between the two hyperoxia paradigms.

### Intrinsic astrocyte hypoxia signaling through HIF2α governs astrocyte abundance

Previous studies have reported conflicting results regarding the function of HIF2α within astrocytes. Duan and colleagues used *GFAP-Cre* mice either to activate the HIF pathway, through deletion of the negative regulator PHD2, or to suppress the pathway through deletion of HIF2α (Duan and Fong, 2019; Duan et al., 2014). They found that pathway activation increases astrocyte density while suppression reduces density, suggesting a possible role for the HIF pathway in regulating astrocyte proliferation. On the other hand, another study using a different *GFAP-Cre* mouse failed to find any effect of astrocytic Hif2α deletion (Weidemann et al., 2010). One possible explanation for this discrepancy is that the efficacy of the *GFAP-Cre-*mediated gene deletion might be variable. For this reason, we decided to monitor HIF pathway activity in our AC-Hif2α-KO mice by staining for VEGF-A. This molecule was an appropriate choice as a readout of HIF signaling, for two reasons. First, VEGF-A is a canonical direct transcriptional target of HIF transcription factors (Pagès and Pouysségur, 2005). Second, in the retina, VEGF-A is the key HIF effector mediating RNFL angiogenesis (Gerhardt et al., 2003; Rattner et al., 2019). Therefore, by staining for VEGF-A, it was possible to probe the aspects of HIF signaling most relevant to retinal vascular development. Using this approach, we validated that the *GFAP-Cre* line used in the Duan et al. studies (Zhuo et al., 2001) does indeed abrogate HIF function. This was shown by the absence of VEGF-A expression from most AC-Hif2α-KO astrocytes at P2, and the absence of astrocytic VEGF-A in at least a subset of mutants at P10.

The P10 VEGF-low mutants exhibited the same phenotype reported by Duan et al. (2014) – i.e. a major decrease in astrocyte abundance and failure of retinal angiogenesis. The astrocyte abundance phenotype is likely due at least in part to proliferation defects, as shown by our Ki67 analysis. Moreover, exposing these mutants to a strong pro-mitotic stimulus – i.e. relative hypoxia via the NOIR paradigm – failed to expand the astrocyte population, further supporting the notion that astrocyte proliferation requires HIF signaling. Our replication of the vascular phenotype is in accordance with recent work – again using the same Cre line – showing that astrocyte-derived VEGF-A is essential for initiation of angiogenesis (Rattner et al., 2019). Overall these results support the notion that astrocyte HIF2α is required both for 1) retinal angiogenesis via induction of VEGF; and 2) hypoxia-induced astrocyte proliferation, in the normal and pathological contexts. The mechanism by which HIF2α promotes proliferation will be an interesting topic for future studies.

Our VEGF-A staining also revealed a subset of VEGF-high AC-Hif2α-KO mice, in which cells retaining HIF function had apparently taken over the astrocyte population. We suggest that this likely occurred as a result of animal-to-animal variability in *GFAP-Cre* recombination efficiency. Cre^−^ astrocytes retaining HIF function are more proliferative than astrocytes with successful HIF2α knockout (Fig. 7B). Therefore, if the Cre^−^ population is large enough, it is conceivable that this competitive advantage could allow wild-type cells to colonize the entire retina. These results point to a potential source of confusion for past studies of HIF and VEGF-A in astrocytes, and highlight the need to validate knockout efficacy on a cell-by-cell basis when genes affecting proliferation are targeted.

### Neonatal hyperoxia alters the trajectory of retinal vascular development

In the NOIR paradigm, vascular development was disrupted in two distinct ways. First, consistent with previous reports (Claxton and Fruttiger, 2003; Morita et al., 2016; West et al., 2005), we found that hyperoxia from P0 prevents the initiation of retinal angiogenesis. Only after return to normoxia do retinal vessels sprout from the ONH. Meanwhile, the embryonic hyaloid vascular system, which normally regresses in conjunction with elaboration of the intrinsic retinal vessels, persisted and in some cases even colonized the neural retina (e.g. Fig. 2D, periphery). Starting hyperoxia at P0 has qualitatively different effects compared to later ages. Hyperoxia from P2 or P4 – ages in the mouse at which retinal vessels have sprouted from the ONH but have not reached the retinal periphery – does not arrest peripheral extension, but instead causes central vaso-obliteration similar to P7-12 OIR (Smith et al., 1994). This observation suggests that initiation of retinal angiogenesis is a distinct and segregable process from peripheral extension of intraretinal vessels, and that each process responds differently to oxygen.

Second, when retinal angiogenesis does eventually begin following return to room air, it does not progress normally. NOIR retinas lacked normal capillary meshwork organization, with severe cases instead displaying dense sheets of endothelial cells without tubular structures. A primary vascular plexus was completed with variable delay, but with long-lasting retinal abnormalities including preretinal vascular membranes; retinal folds suggestive of tractional retinal detachments; reactive Müller glia; and vitreous hemorrhage. These findings differ from a recent study that exposed CD-1 mice to P0-4 hyperoxia, in which the effects on retinal vasculature were reported to be mild and transient (Morita et al., 2016). The reason for the discrepancy between the two studies is unclear, as key aspects of study design – including the O_2_ percentage used – were similar. Since we observed persistent pathology only in ∼50% of treated animals, one possibility is that random chance played a role in the severity of effects observed in the previous study. By contrast, our results are consistent with pathologies seen in young adult mice exposed to hyperoxia from P0-P7 (McMenamin et al., 2016). Together these findings suggest that initiation of hyperoxia earlier than the traditional OIR model (Smith et al., 1994) can produce a different vascular phenotype, with the vitreoretinopathy and tractional detachments potentially mirroring some aspects of ROP (Foos, 1987) that are not well modelled by OIR.

### Role of astrocytes in oxygen-induced vasculopathy

Our results from the NOIR paradigm show a strong correlation between the magnitude of astrocyte overproduction and the severity of the vascular delay and disarray. Since astrocyte phenotypes precede vessel growth and occur in peripheral retina prior to the arrival of vessels, we favor a model in which hyperoxia affects astrocytes, and astrocytes in turn contribute to pathologic retinal vascularization. This would be consistent with other manipulations of astrocyte population size during development, each of which demonstrate that vascular development is highly sensitive to the number of astrocytes within the angiogenic template (Duan and Fong, 2019; Fruttiger et al., 1996; O’Sullivan et al., 2017; Puñal et al., 2019; Tao and Zhang, 2016). However, our experiments cannot exclude the possibility of additional factors explaining the correlation between astrocytic and vascular phenotypes. Furthermore, additional astrocyte abnormalities may arise as a result of vascular development errors, given that astrocytes depend on signals from endothelial cells for their own maturation (Mi et al., 2001; Sakimoto et al., 2012; Selvam et al., 2018). Further work will be required to better elucidate the mechanisms by which astrocytes and endothelial cells interact during development and disease.

### Implications for ROP

Certain defining features of ROP, such as delayed peripheral vascularization, astrocyte hyperproliferation, and long-lasting retinopathy, have been challenging to model in the mouse and are absent in the classic P7-P12 mouse OIR model (Gariano, 2010). Therefore, while traditional mouse OIR remains an extremely useful disease model, the neontatal hyperoxia paradigm we employed here has several features that make it a useful complement to traditional OIR. First, the astrocyte hyperproliferation phenotype we describe in this study could provide useful insight into formation of the astrocyte-dense fibrovascular ridge that forms in the mid-periphery of ROP retinas (Chen et al., 2018; Foos, 1987; Sun et al., 2010). Second, the NOIR paradigm could aid in understanding ROP variants that arise at earlier stages of retinal development than those modelled by the traditional OIR model. As improvements in neonatology improve survival rates for the most premature infants, models of these early retinopathies may become increasingly important and relevant to clinical practice. Finally, the enduring retinopathy produced by neonatal hyperoxia – rather than the spontaneously resolving vaso-obliteration and neovascularization in OIR – could provide valuable developmental outcome measures for potential therapies.

Current treatment strategies for ROP target Phase II and aim to reduce neovascularization either by 1) ablating ischemic peripheral retina to eliminate the source of abnormal proangiogenic signals or 2) blocking VEGF signaling. While overall helpful in preserving functional vision in affected children, both approaches intervene relatively late in the pathogenesis of ROP. Additionally, laser and cryoablation are inherently destructive procedures, and concerns remain regarding possible ocular and systemic side effects of anti-VEGF therapy during development. Better understanding of the pathogenesis of ROP and the role of astrocytes could reveal new targets for treating or preventing retinal vascular disorders. Our results suggest that inhibiting astrocyte overproliferation could be a promising therapeutic avenue for preventing vessel pathology. In our experiments we were unable to assess whether suppressing hypoxia-induced proliferation improved vessel phenotypes, because AC-Hif2a-KO mutants lacked all intrinsic retinal vasculature. This was likely due to the absence of VEGF-A expression from astrocytes, without which retinal angiogenesis cannot initiate (Rattner et al., 2019). To critically test whether astrocyte proliferation is a promising therapeutic target, it will be necessary to identify additional and more selective molecular determinants of hypoxia-induced proliferation.

Our study sets the stage for identifying such factors and for testing their role in vascular pathology. With the introduction of the NOIR paradigm, we provide an experimental model for addressing the mechanisms of hypoxia-induced astrocyte proliferation and their consequences for retinal pathology. We propose that this approach will ultimately facilitate identification of molecular pathways that suppress astrocyte proliferation without interfering with angiogenesis. Further, once such pathways are identified, it will be possible to test the potential of these pathways as targets to ameliorate developmental vascular pathology.

## Acknowledgements

This work was supported by the National Eye Institute (EY030611 to J.N.K; EY5722 to Duke University); the Ruth K. Broad Foundation (J.N.K.); Research to Prevent Blindness (Unrestricted Grant to Duke University); and a Duke University Holland-Trice award (J.N.K.). We thank Ari Pereira for mouse husbandry; William Marcus for technical assistance; Xi Chen and Joseph Brzezinski for comments on the manuscript; and William Spencer for help with photography. The funders had no role in study design, data collection and analysis, decision to publish, or preparation of the manuscript.

## Supplemental Figures and Legends

**Figure S1.**
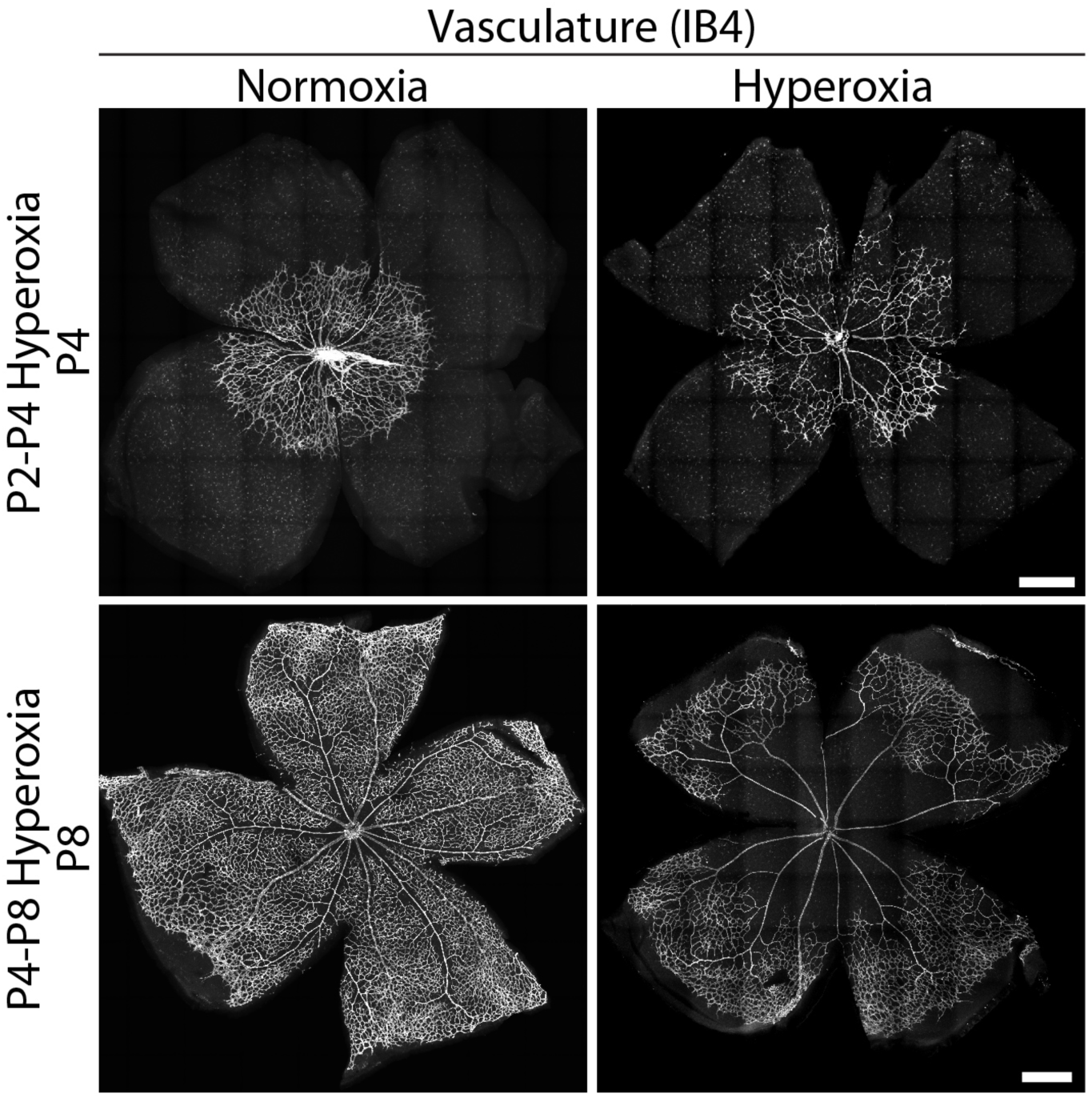
Effects of starting hyperoxia at P2 or P4 on vascular development. Representative whole-mount images of retinal vasculature from animals reared in 75% O_2_ from P2-P4 (top) or P4-P8 (bottom). Progression of vascular wavefront is similar to normoxic littermate controls in both treatment paradigms. However, central vaso-obliteration is observed in treated animals, similar to standard P7-P12 OIR paradigm.

**Figure S2.**
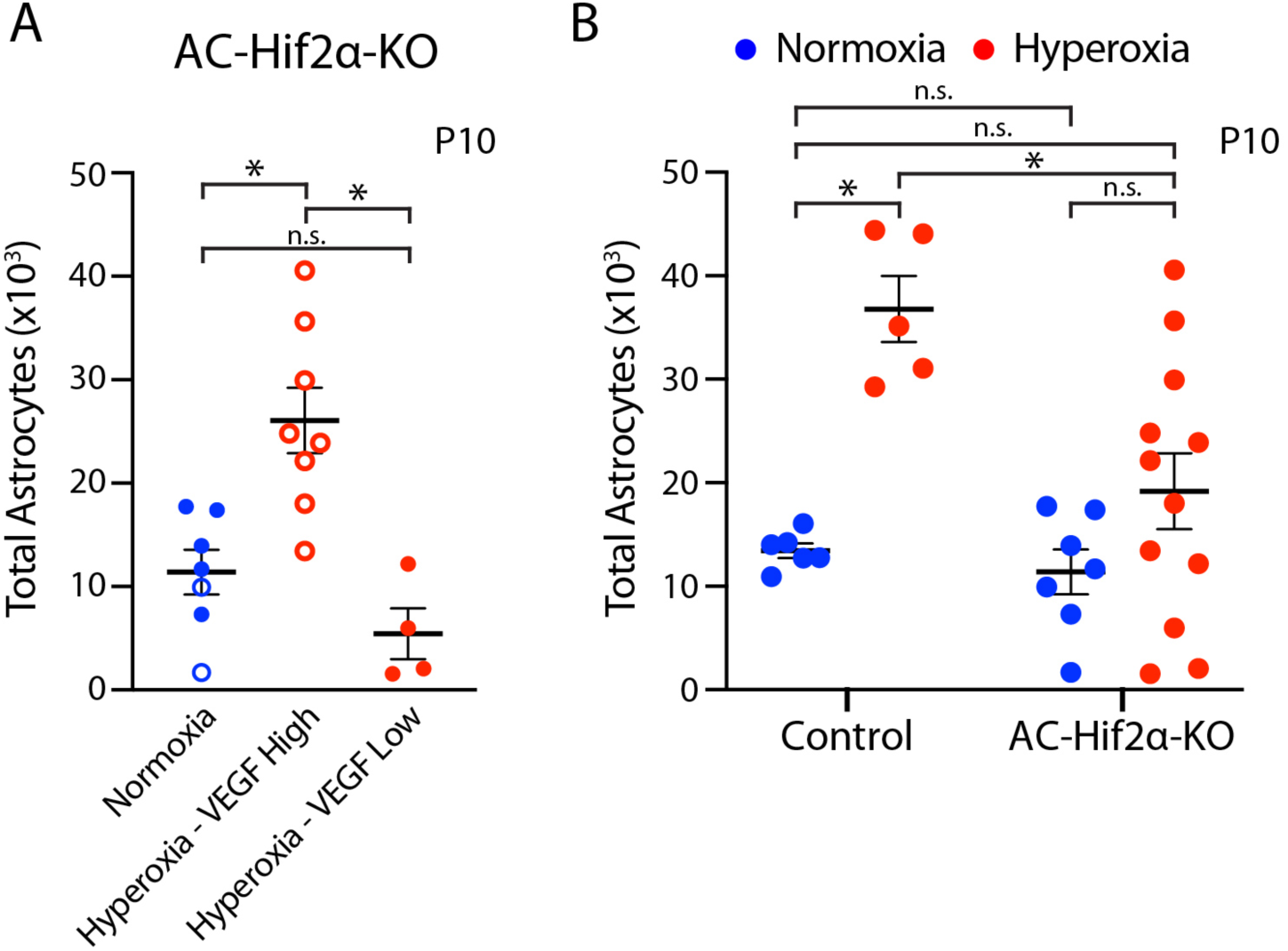
Further characterization of VEGF-High and VEGF-Low classes of Hif2α mutants. **A**) VEGF-high and VEGF-low mutants (P10) have significantly different astrocyte production responses in the NOIR paradigm. This finding provides further evidence that the two classes of mutants are fundamentally different and should be considered different experimental groups. Normoxia group includes both VEGF-low (open circles) and VEGF-high (closed circles) KO mice. One-way ANOVA, F (2,16) = 13.5, p = 0.0004. Asterisks denote significant differences by Tukey’s post-hoc test. VEGF-high vs. VEGF-low, p = 0.0007. Normoxia vs. VEGF-high, p = 0.0032. Normoxia vs. VEGF-low, p = 0.4079. **B**) Total astrocyte numbers at P10 in mutants and littermate controls. Graph is the same as Fig. 8C, except that all AC-Hif2α-KO mutants have been plotted regardless of VEGF status. Even when no distinction is made between mutants based on VEGF expression, there is still a significant effect of genotype on astrocyte number (Two-way ANOVA, main effect of genotype F(1,26) = 7.780, p = 0.0098 ; main effect of oxygen F(1,26) = 19.49, p = 0.0002 ; interaction, F(1,26) = 4.846, p = 0.0368). Furthermore, a post-hoc test (Holm-Sidak) shows that loss of Hif2α significantly blunts the effect of hyperoxia even when VEGF-high animals are included in the analysis (control hyperoxia vs. mutant hyperoxia, p = 0.0051). Consistent with this finding, we did not detect a significant astrocyte number difference between normoxic and hyperoxic mutants (Holm-Sidak, p = 0.2337). Therefore, even though there is strong biological justification for excluding VEGF-high mutants from the experiment, failing to do so does not change our central conclusion that Hif2α is required for the astrocyte proliferative response in the NOIR paradigm. Other P values noted on graph: normoxia controls vs. normoxia mutants, p = 0.6874; normoxia controls vs. hyperoxia mutants, p = 0.3947. Error bars, mean ± s. d.

